# Neuroplasticity of directed connectivity in long-term meditation: Evidence from EEG Granger causality

**DOI:** 10.1101/2025.07.01.662528

**Authors:** Vasil Kolev, Kosio Beshkov, Peter Malinowski, Antonino Raffone, Juliana Yordanova

## Abstract

The objective of the present study was to characterize the effects of long-term meditation (LTM) on directed connectivity patterns during resting-state and meditative brain states. Specifically, it was aimed to identify major cortical sources of information flow and target regions of information influx, and to reveal which frequency-specific oscillatory networks are critically involved in directing information flux in experienced meditators.

Multivariate Granger causality (GC) was computed from high-resolution EEG signals recorded from long-term (LTM, n = 22) and short-term meditators (STM, n = 17) in four conditions: rest, Focused Attention Meditation, Open Monitoring Meditation, and Loving Kindness Meditation. GC was analyzed in the time and frequency domains to assess frequency-specific networks supporting the directed connectivity between key cortical regions (frontal and parietal) in the two hemispheres.

According to the results, long-practice meditation was characterized by a significant increase of information flow (1) from posterior to frontal cortical regions, and (2) across frontal regions of the two hemispheres. These dominant transfers were supported by multi-spectral oscillatory networks involving theta, alpha and beta frequency bands, with most prominent expression of GC alpha peak. This pattern of enhanced information transfer in LTM relative to STM was observed in both resting state and each meditation state.

These results suggest that long-term meditation is associated with a shift in resting-state brain dynamics toward reduced reliance on slow, undirected intrinsic oscillations, and enhanced directional connectivity in frequencies linked to attention and cognitive control. The dominant posterior-to-anterior directionality points to a reorganization of cognitive control networks that may support the phenomenological qualities of extensive meditation (sustained attention, internal attention, present-moment awareness, and reduced cognitive elaboration). The similarity of between-group differences in directionality patterns across states points to a neuroplastic effect of long-term meditation and highlights meditation as a potential model for investigating adaptive neuroplasticity in large-scale brain networks.

## 1. NTRODUCTION

Meditation and mindfulness have been integrated into therapeutic interventions in both clinical conditions such as stress, chronic pain, depression and anxiety (Van Aalderen et al., 2012; Davis and Hayes, 2011; Goyal et al., 2014; Gu et al., 2015; Hofmann et al., 2011) and non-clinical contexts such as workplaces, education, sports, and criminal justice systems (Heckenberg et al., 2018; McKeering and Hwang, 2019). Accordingly, scientific research of various forms of contemplative practice, including meditation, has gained increasing interest (Ospina et al., 2007; Chiesa, 2009, 2010; Rubia, 2009; Chiesa and Serretti, 2010; Green and Turner, 2010). One goal of this research is to provide objective scientific explanations of the therapeutic effects of meditation interventions. A larger scientific goal is to understand the neural grounds of meditation states in terms of mechanisms and networks that support them.

One established approach to explore the neural correlates of meditation is to study highly experienced meditators who have acquired mental skills over many years and are able to maintain different meditative states at will (Carter et al., 2005; Lutz et al., 2004; Manna et al., 2010; Marzetti et al., 2014). As meditation experience accumulates, the ability of achieving and sustaining specific meditative states substantially increases thus enhancing the reliability of neural measurements. Also, the repeated engagement of various cognitive systems and associated brain networks after long practice is expected to induce neuroplastic changes that would be more clearly delineated and identified as mediators of positive outcomes (Brefczynski-Lewis et al., 2007; Pace et al., 2009; Baron Short et al., 2010; Manna et al., 2010; Yordanova et al., 2021). Thus, studying highly experienced practitioners offers a unique opportunity to extract and explore reliably the neurofunctional substrates of meditative states.

Previous studies of experienced meditators have indicated that extensive meditation practice is associated with modulations and neuroplasticity of major cognitive networks (Raffone et al., 2019; Malinowski, 2013; Tang et al., 2015; Lutz et al., 2015; Raffone and Srinivasan, 2017), with large-scale integrative mechanisms playing a crucial role for maintaining conscious brain states (Dehaene et al., 1998, 2015; Raffone et al., 2014; Tononi and Edelman, 1998; Tononi et al., 1992; Varela et al., 2001). Meditation may not only involve process-specific brain networks, but may also train integrated brain states supporting a relevant interplay with the autonomic nervous system (Malinowski and Shalamanova, 2017; Tang and Posner, 2014). In the framework of large-scale networks interaction (Bressler and Menon, 2010), measures of brain connectivity have been identified as plausible markers of meditative states.

Currently, a variety of different approaches are used to measure brain connectivity (rev. Mueller et al., 2016). They are essentially based on signals derived from functional magnetic resonance imaging (fMRI) or electroencephalogram (EEG). Such approaches typically include analyses of coherence, correlation, autocorrelation, and phase-synchrony to assess the coherent and synchronous involvement of distant cortical or sub-cortical regions representing key nodes of relevant networks (functional connectivity). Previous studies of functional connectivity have provided evidence for meditation-related alterations of the connections between and within major neurocognitive networks verifying their involvement in the maintenance of meditative states.

For example, using fMRI Hasenkamp et al. (2012; Hasenkamp and Barsalou, 2012) have demonstrated that even at rest, greater meditation experience in focused attention meditation (FAM) is associated with increased functional connectivity among attentional networks as well as altered connections between anterior and posterior elements of the Default Mode Network (DMN). Likewise, Barros-Loscertales et al. (2021) have found that long-term meditation practice increases functional connectivity within attentional and cognitive control networks and decreases connectivity between these networks and the DMN. EEG studies of functional connectivity in the frequency domain have shown that compared to novice meditators, highly experienced meditators exhibit a strong theta synchronization of both attentional and executive networks in left parietal regions during resting state as well as other meditation states, and only the connectivity of lateralized beta networks differentiate meditation styles (Yordanova et al., 2021). Using EEG coherence has further demonstrated that state-unspecific connectivity patterns from theta and alpha bands characterize network organization in experienced meditators (Yordanova et al., 2020). Together, these observations imply that neuroplasticity of frequency-specific networks supporting inter-regional connections may underpin the emergence of unique meditation states in expert meditators.

It is acknowledged however that cortical connectivity can be more deeply understood by analyzing directed information flow, in addition to assessing the synchronous involvement of distant brain regions (Hillebrand et al., 2016; Pullon et al., 2020). Measures of functional connectivity reflect regional synchrony, but they cannot capture the direction of information flow. Therefore, a number of directed measures have been developed to understand inter-regional information flux (“directed functional” or “effective” connectivity). In general, they quantify whether the information from a distant brain region has occurred just before that of the target region and whether it has influenced the fMRI/EEG time series at the target region (Blinowska et al., 2004; Mueller et al., 2016). The application of one such established measure – Granger causality (GC) – has revealed that directed and undirected measures of connectivity may provide independent and complementary information about dynamic changes of brain states (Pullon et al., 2020).

Despite the potential of effective connectivity to refine the characterization of meditation states, studies on directed information flow in meditation remain scarce. Rao et al. (2018) computed Granger causality from fMRI signals and to reported a reduction of information outputs from insula, anterior cingulate and orbitofrontal cortices to limbic structures during meditation and rest. However, the frequency specificity of networks underpinning directed information flow in long-term meditation has not been explored although frequency may be a most relevant signature of meditative states (Lutz et al., 2008; Marzetti et al., 2014; Yordanova et al., 2020, 2021; Guidotti et al., 2023). Also, based on the Global Neuronal Workspace Model of consciousness, (Dehaene et al., 2015), it is plausible to suggest that the complexity of conscious states in meditation is supported by free information flux between cortical regions engaging frequency-specific networks.

The objective of the present study was to complement the characterization of functional connectivity patterns in highly experienced meditators by EEG-based analysis of directed information flow. Specifically, it was aimed (1) to explore if long-term meditation practice affects the direction and amount of information streams in the brain, (2) to highlight major cortical sources of information influx and outflux in experienced meditators, and (3) to reveal which frequency-specific oscillatory networks are critically involved in directed information flow during rest as well as maintenance of different meditative states. For that aim, multivariate Granger causality was computed and analyzed in experienced and novice meditators in four conditions: REST, Focused Attention Meditation (FAM), Open Monitoring Meditation (OMM) and Loving Kindness Meditation (LKM) (Yordanova et al., 2021).

## 2. MATERIALS AND METHODS

### Participants

A previously reported EEG data set was used for the present new analyses (Yordanova et al., 2021). The group of long-term meditators (LTM) consisted of twenty-two healthy volunteers (mean age = 44.2 years, age range 26-70 years, 4 females). They were right-handed and did not report any history of movement disorders, neurological, psychiatric or somatic diseases. These participants were monks, nuns and novice practitioners residing at Amaravati Buddhist Monastery, in Southern England, and at Santacittarama Monastery, in Central Italy. Practices at both monasteries are aligned with the Tai Forest Theravada Buddhist tradition which is now established, widely acknowledged and influential in the West. Participants practiced FAM (Śamatha), OMM (Vipassanā) and LKM (Metta) meditation forms in a balanced way, often in integrated sessions, including silent meditation retreats (3 months per year). Meditation expertise is measured in hours taking into account both practice in the monastic tradition and practice before monastic life. In this tradition, the monks, nuns and novice practitioners typically practice two hours per day with the monastery community, with a regular intensification of practice during retreats. As suggested by the abbots of the monasteries, we estimated participants with an average of 100 hours of practice per month during monastic life, with a balance of FAM, OMM and LKM facets of meditation. The lifetime duration of meditation practice of the participants was estimated as a mean value = 19358 hours (SE=3164), range 900-50600 hours.

Another group of 17 participants (mean age = 45.5, age range 32-58 years, 9 females) with less than 250 h of meditation experience in secular mindfulness or Buddhist traditions also was studied. These short-term meditators (STM) further practiced the instructions of FAM, OMM and LKM for 10 days before the study, 20 min per day for each form of meditation. They met the same criteria: right-handedness and lack of any somatic, neurologic or psychiatric diseases.

The study had prior approval by the dedicated Research Ethics Committee at Sapienza University of Rome, Italy. All participants gave informed consent before participation according to the Declaration of Helsinki.

### Experimental design

Participants had to perform a non-meditative rest condition and three meditation conditions: FAM, OMM and LKM. The switching between conditions was cued by voice. The instructions for the four conditions, which were written together with the abbot of Amaravati Monastery, the internationally recognized teacher Ajahn Amaro, were as follows: Rest: “Rest in a non-meditative relaxed state, without falling in sleep, while allowing any spontaneous thoughts and feelings to arise and unfold in the field of experience”. Focused attention (Śamatha) meditation (FAM): “Sustain the focus of attention on breath sensations, such as at the nostrils, noticing readily and with acceptance any arising distraction, such as on thoughts or stimuli, and in case of detected distraction, return readily and gently to focus attention on the breath sensations”. Open monitoring (awareness) meditation (OMM): “With an open receptive awareness, observe the contents of experience as they arise, change and fade from moment to moment, without restrictions or judgments – such contents including breath and body sensations, sensations arising from contact with external stimuli, feelings and thoughts”. Loving kindness (Metta) meditation (LKM): “Generate and sustain Metta, acceptance and friendliness towards yourself and the experience in the present moment, as well as towards any being, in any state or condition”.

### Procedure

#### EEG recordings

EEG was recorded with eyes closed in blocks of duration of approximately 2.5-3 minutes each, while the above mentioned conditions were repeated two times. In such a way, four blocks were repeated twice in the following order: REST, FAM, OMM, LKM. Thus, there were approximately 5-6 minutes (2 blocks x 2.5-3minutes) of total recording time for each condition. EEG was acquired by a mobile wireless system Cognionics (https://www.cognionics.net/mobile-128) using an electrode cap with 64 active Ag/AgCl electrodes located in accordance with the extended international 10/20 system and referenced to linked mastoids. Electrode impedances were kept below 10 kOhm. EEG signals were collected at a sampling rate of 500 Hz. For control of ocular artefacts, vertical and horizontal electro-oculogram (EOG) was also recorded. For experienced meditators, the experiment was conducted at the two monasteries in a quiet, dark room suitable for meditation and recording EEG. All participants were tested with the same experimental design.

#### EEG pre-processing

EEG pre-processing was performed by means of Brain Vision Analyzer 2.3 (Brain Products GmbH, Germany). Granger causality analysis of EEG was conducted by means of software developed on Matlab R2013b (The Math Works Inc.).

EEG traces were visually inspected to reject epochs with noise or non-physiological artefacts. Bad channels were interpolated according to Hjorth (1975). All EEG traces were EOG corrected by means of independent component analysis (ICA, Makeig et al., 1997). EEG data were down-sampled to reduce off-line the recording frequency of 500 Hz to 250 Hz for data analysis.

To achieve a reference-free evaluation, all data analyses were performed after current source density (CSD) transform of the signals (Perrin et al., 1989; Nunez et al., 1997). The exact mathematical procedure is presented in detail in Perrin et al. (1989). The algorithm applies the spherical Laplace operator to the potential distribution on the surface of the head. The CSD transform replaces the potential at each electrode with the current source density of the electrical field calculated from all neighbor electrodes, thus eliminating the reference potential. When applied with dense electrode arrays (48–256 electrodes, 64 in the present study), this procedure provides excellent estimates of the bioelectric activity of the cortical surface (Nunez and Pilgreen, 1991). In the present analyses, to improve spatial resolution and reduce the volume conduction between electrodes, CSD was applied following with the parameters: order of splines = 10, maximal degree of Legendre polynomials = 4, lambda = 1E^-7^. Edge electrodes were excluded from all analyses, so that the number of channels was reduced. All analyses were carried out with CSD transformed data from 50 electrodes.

EEG recordings from all conditions were segmented in equal-sized non-overlapping epochs of 4.096-s duration. The average number of epochs for each condition/participant was 58 (±20). There were no statistically significant differences in the number of accepted trials between the two groups for each condition (REST, FAM, OMM, LKM).

#### Analysis of Granger causality (GC)

To support the applied methodology, the formalism of Granger causality in the time and frequency domains is briefly presented in the following section.

#### Time domain formulation

Granger causality (Granger, 1969) is a method for the estimation of “Granger causal” relations between time series. Granger causality tries to see if the history of a time series Y holds information about the future of a different time series X which is not present in the history of X itself. In order to calculate this, two model equations are constructed:

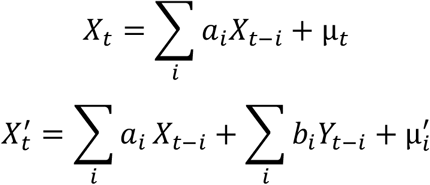

The *μ* terms are independently and identically distributed (iid) residuals and we can denote their variance as *var*(*μ*) = Γ and *var*(*μ*^′^) = Γ′. Then one can define the Granger causality statistic from Y to X as

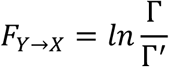

In more general situations when there are more than two signals, a choice has to be made whether to claim that there is a Granger causal influence from Y to X if Y improves the prediction of X on its own *or* in combination with all the other variables. The former case is known as *unconditional Granger causality* and has the disadvantage that it leads to spurious causalities because it does not take into account higher order interactions between variables. However, this method is sometimes preferable as it is less strict and pairwise dependencies are still interpretable. The second method is known as *conditional Granger causality* and for each connection, it takes into account all other contributing sources. This approach is much stricter and it correctly reflects the higher order structure of connectivity. It can be expressed formally with the following equations:

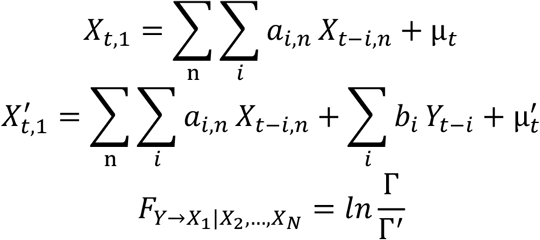

These are essentially the same equations as presented above except for the fact that a sum is added that goes over all other variables except Y.

#### Frequency domain (spectral) formulation

One of the main reasons that Granger causality has generated so much interest in neuroscience is the fact that it can be reformulated to estimate the connectivity between neural sources in specific frequency bands (Geweke, 1982, 1984).

In order to calculate spectral causality it is not correct to apply band-pass filters to the data and then apply Granger causality. Instead, a Fourier transform is applied to the time lagged regression coefficients and then the frequency specific Granger causality is calculated according to the formula:

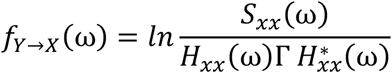

Here *S* represents the cross-power spectral density, *H* is the transformed (following Geweke, 1982) transfer function, with the star denoting the complex conjugate, and Γ is the residual covariance matrix. The full derivation of the frequency domain Granger causality is presented in detail in Chen et al. (2006) and Barnett and Seth (2014).

### Present analysis of GC

#### GC computation pre-processing

Granger causality was analyzed using the publicly available MATLAB package (MVGC Matlab toolbox, Barnett and Seth, 2014). We paid particular attention to pre-processing steps given the sensitivity of GC to standard manipulations (Bressler and Seth, 2011; Seth, 2010). Analysis of Granger causality requires the analyzed signals to be stationary and not collinear (Seth, 2010). To avoid problems associated with these properties, the following additional pre-processing steps were undertaken.

First, as outlined before, a Laplacian filter (CSD) was applied to remove correlations in nearby electrodes, thereby also reducing the collinearity between signals.

Second, following the requirements (Barnett and Seth, 2011, 2014), to remove mains-electricity line-noise (50 Hz) as well as its harmonic at 100 Hz which may lead to nonstationarity, notch filters (49–51 Hz and 99–101 Hz) were applied to the raw data. No other filtering was carried out. Also, since the length of the initially selected epochs in the data (4.096 s, 250 Hz) has been demonstrated to exhibit nonstationary features (e.g., Barrett et al., 2012), these epochs were additionally processed. They were divided into stationary non-overlapping segments of 1.024-s length. This length also was chosen to approach a balance between stationarity (shorter time series are more likely to be stationary) and model fit (longer time series support better parameter estimation for locally valid linear autoregressive models). This segmentation increased reliability by leading to a mean of 232±40 artefact-free EEG epochs per subject/per condition. After this step, the GC estimation still took an intractable amount of time. To ensure a reasonable model order for autoregressive modelling (Seth, 2010; Brovelli et al., 2004) and a reasonable amount of computation time, the selected artefact-free epochs were further subsampled by taking every fourth point from each time series. This effectively reduced the sampling rate by a factor of four (62.5 Hz). In this way, each epoch used for final analyses had a length of 1.024 s comprising 64 data points.

The third pre-processing step was to compute sensor-based regions of interest or electrode clusters (C) covering the left and right anterior and posterior cortical regions, for which specific connectivity patterns in meditation have been reported (Yordanova et al., 2020). We performed clustering as demonstrated in Fig. 1. The clustering scheme involved averaging the signals from the following electrodes: C1 (left anterior) = [F7, F5, F3, F1, FC5, FC3, FC1, C5, C3, C1]; C2 (right anterior) = [F8, F6, F4, F2, FC6, FC4, FC2, C6, C4, C2]; C3 (left posterior) = [CP5, CP3, CP1, P7, P5, P3, P1, PO7, PO3, O1]; C4 (right posterior) = [CP6, CP4, CP2, P8, P6, P4, P2, PO8, PO4, O2]. After the spatially averaged single-trial EEG of each defined cluster were computed, to avoid further confounds due to nonstationarity, a linear detrend was applied (Bernasconi and Konig, 1999; Hesse et al., 2003). Single-trial time-series corresponding to C1, C2, C3 and C4 were extracted for each subject and each condition and were used in subsequent analyses.

**Figure 1.**
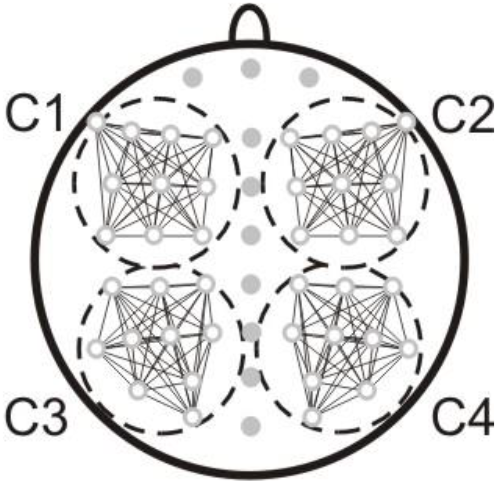
Schematic presentation of topographic clusters used for analysis: C1 (left anterior) = [F7, F5, F3, F1, FC5, FC3, FC1, C5, C3, C1]; C2 (right anterior) = [F8, F6, F4, F2, FC6, FC4, FC2, C6, C4, C2]; C3 (left posterior) = [CP5, CP3, CP1, P7, P5, P3, P1, PO7, PO3, O1]; C4 (right posterior) = [CP6, CP4, CP2, P8, P6, P4, P2, PO8, PO4, O2].

#### Computation of GC

To compute GC, the openly available MATLAB package MVGC Matlab toolbox (Barnett and Seth, 2014) was first used to estimate the model order. For each single EEG epoch of the data, the recommended model order was computed as defined by the Akaike information criterion (AIC, Akaike, 1974; McQuarrie and Tsai, 1998). The 95th percentile of the values obtained was 10 (corresponding to 160 ms) and was used as model order throughout the GC analysis. The need of model order selection is to balance the number of parameters (as determined by the maximum order - lag) so as to achieve the best model fit to data while avoiding overfitting a finite data sequence. The next step was to maximize the likelihood function for the respective models. The MVGC toolbox facilities were used to obviate the need to estimate the reduced model parameters separately from the data. Specifically, to yield estimates asymptotically equivalent to the maximal likelihood estimate, ordinary least squares (OLS) were used (Hamilton, 1994). The estimated regression coefficients in the time domain were then transformed to an auto-covariance sequence through the Yule-Walker equations (Barnett and Sett, 2014), which was used to compute GC in the time domain. Finally, this sequence was used to calculate the Granger causalities in the frequency domain.

#### Statistical Analysis

##### Time domain

Statistical evaluation of GC in the time domain was performed using a parameter fraction of significant connections (FSC). It reflects the normalized ratio between the number of significant GC connections from all participants in each group (LTM, n = 22, STM, n = 17) and the number of all connections included in the analysis for each group for each of evaluated pair (C1→ C2, C3, C4; C2 → C1, C3, C4; C3 → C1, C2, C4; and C4 → C1, C2, C3). Fraction of significant connections is expressed as the number of epochs*participants with significant connections divided to the total number of epochs*participants. To compute FSC, the following steps were followed: (1) After GC was computed in the time domain for each single trial, subject, condition and between-cluster pair, a statistical test based on the theoretical null distribution (F-distribution) was performed for each trial (https://users.sussex.ac.uk/~lionelb/MVGC/html/mvgc_pval.html). The test determines if the GC of single-trial between-cluster pair is significant at individual level. (2) Significant pairs were coded with 1, and non-significant – with 0. (3) For each between-cluster pair, these values were summed up for all trials of all subjects in each group separately (LTM and STM) and were divided by the sum of all trials from all subjects in the respective group. The so computed normalized FSC was used for visualization and between-group descriptive comparisons.

##### Frequency domain

In order to compare the Granger causalities across frequencies for the LTM and STM groups, the following steps were followed: (1) After computing GC in the frequency domain, first, the Granger spectrum for each subject, each condition, each single trial, and each between-cluster pair was discretized into 1Hz bins. (2) Then, to compare the GC between LTM and STM, all GC values within each 1Hz frequency bin for each trial/condition/pair in each group were concatenated and subjected to a non-parametric pair-wise Kruskal-Wallis test. Since this type of analysis involves multiple comparisons, the results were corrected with a false discovery rate (FDR) correction, as recommended for GC analysis in the frequency domain (Barnett and Seth, 2014).

## 3. RESULTS

### 3.1. Time domain analysis

Main results of the analysis of the fraction of significant connections computed using GC are presented graphically in Figure 2 for the groups of STM and LTM in four conditions: REST, FAM, OMM and LKM.

**Figure 2.**
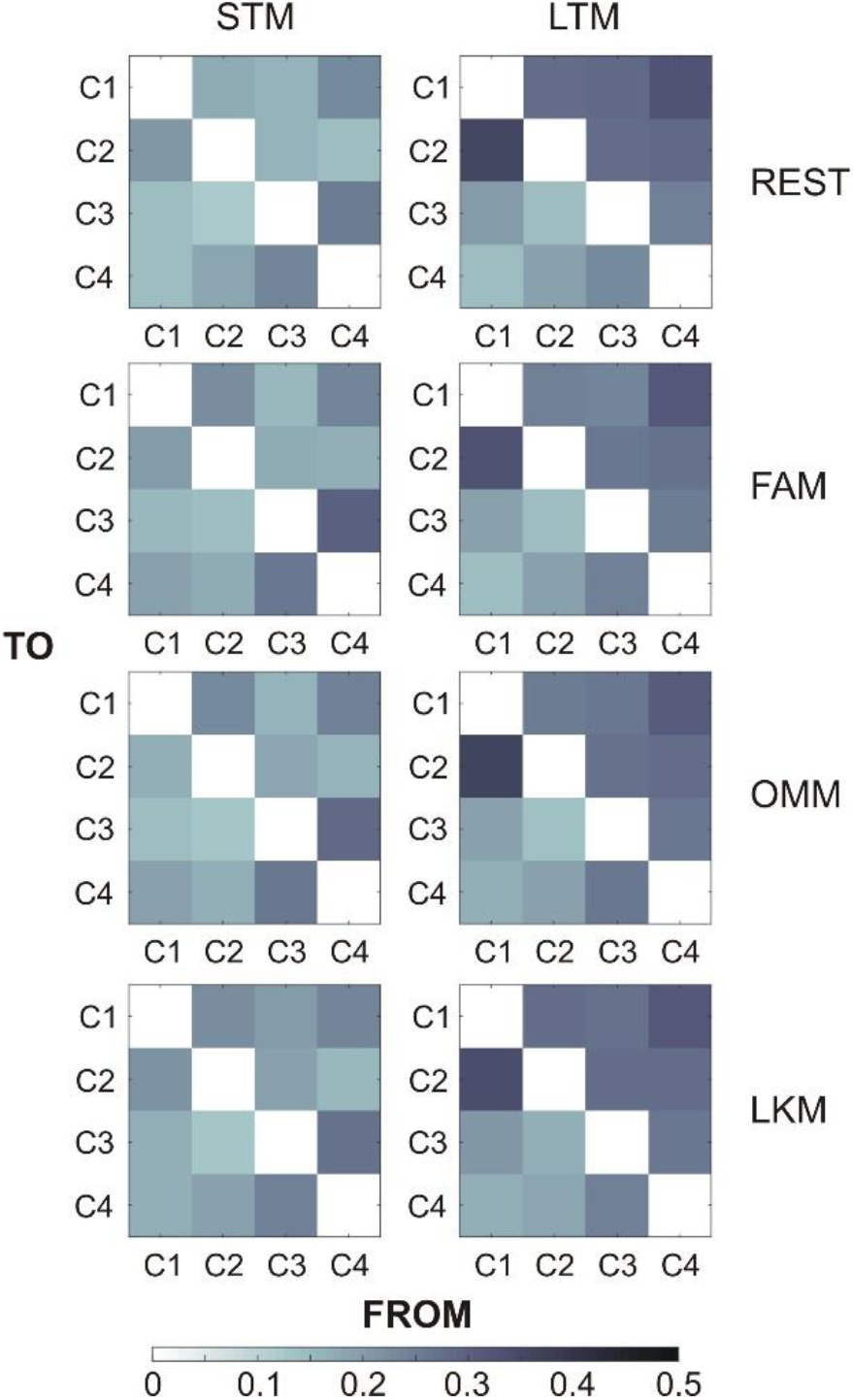
Time-domain analysis of Granger causality. Normalized fraction of significant connections (FCS) yielded after GC computation in the time domain is presented for long-term meditators (LTM – right) and short-term meditators (STM – left) in four conditions REST, Focused Attention Meditation (FAM), Open monitoring Meditation (OMM), and Loving Kindness Meditation (LKM). Horizontal axis: FROM C1, C2, C3, C4. Vertical axis: TO C4, C3, C2, C1.

Figure 2 demonstrates that in each of the four conditions (REST, FAM, OMM, and LKM), the dominant connectivity pattern in STM was characterized by pronounced inter-connections between the two posterior clusters C3 and C4.

In contrast, the connectivity pattern in LTM was dominated by increased information flows across the two frontal clusters C1 and C2, mainly from C1 to C2 as indicated by the higher fraction of significant connections. Figure 2 further reveals an increased information flow from posterior (C3 and C4) to the frontal (C1 and C2) clusters in LTM. Figure 3 depicts schematically the between-group differences for these clusters and is presented here to facilitate comparisons with findings in the frequency domain (see below). These main patterns of directed connectivity were stable across both resting state and meditation conditions in both STM and LTM (Figs. 2 and 3).

**Figure 3.**
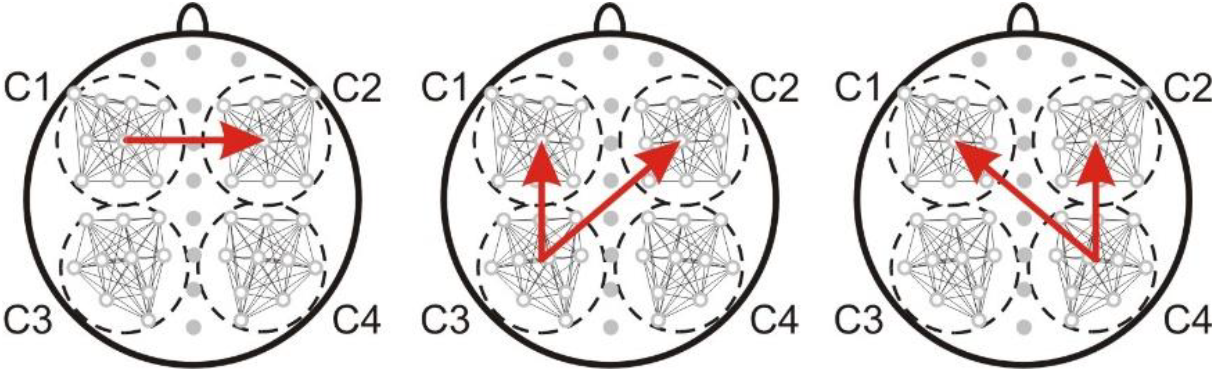
Schematic presentation of observable between-group differences (LTM > STM) in FSC computed from Granger causality in the time domain during REST. The same pattern was observed in meditation conditions (FAM OMM, LKM). Clusters: C1, C2, C3, C4. Arrows indicate the direction of significantly different information flows.

### 3.2. Frequency domain analysis

Frequency-domain representations of GC are demonstrated in Fig. 4. Observation of these representations during rest revealed the presence of a dominating broad peak in LTM with a maximum in the alpha band (8-10 Hz), encompassing frequencies from both lower (theta) and higher (beta) bands. The broad alpha peak in LTM was most prominent for directed transfer connections from posterior (C3 and C4) to anterior (C1 and C2) clusters. It was also present for information transfer across anterior clusters (C1→ C2 and C2 → C1), being however much less expressed in comparison with the posterior-to-anterior transfers.

**Figure 4.**
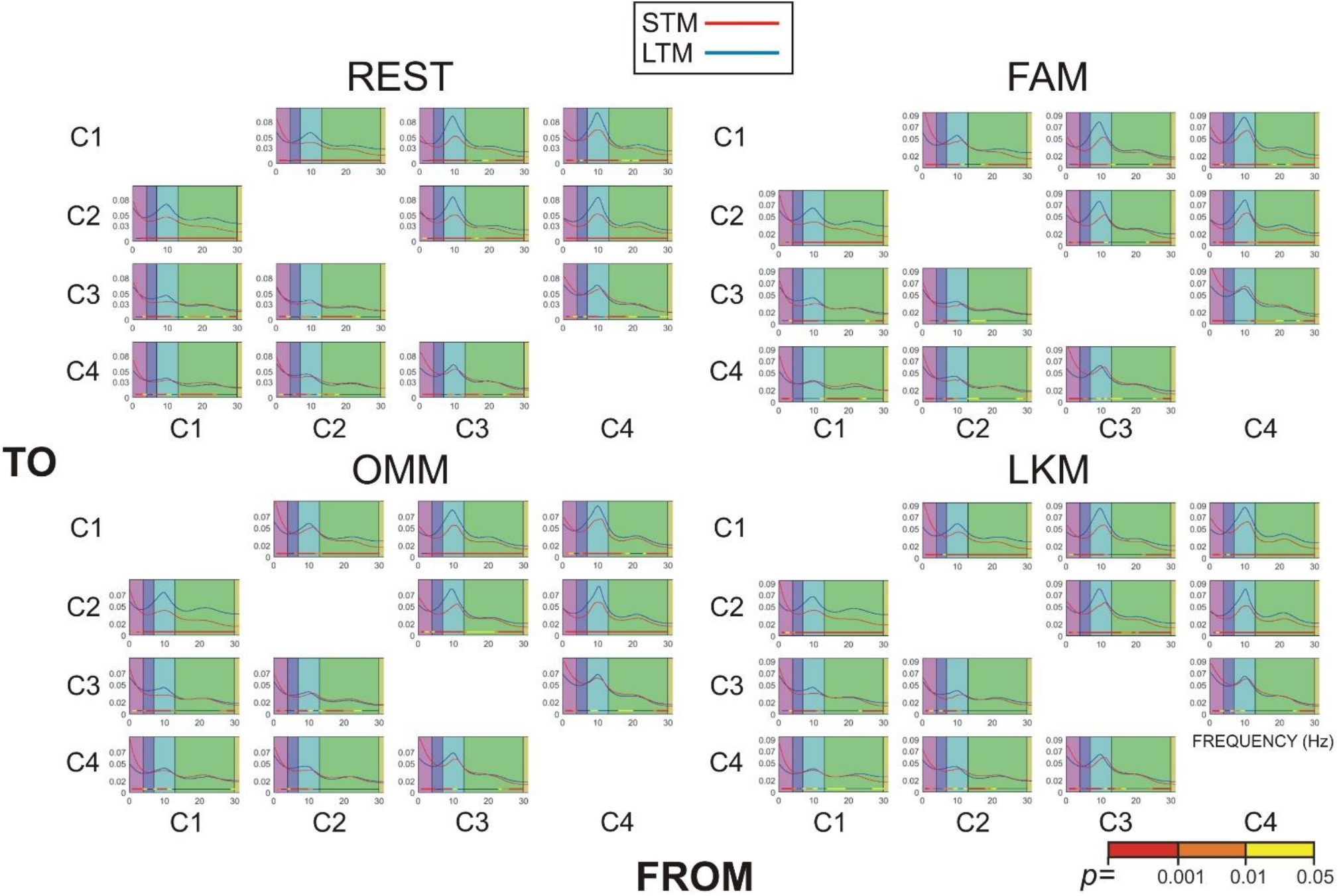
Frequency representation of Granger Causality in long-term meditators (LTM – blue line) and short-term meditators (STM – red line) in four conditions REST, Focused Attention Meditation (FAM), Open monitoring Meditation (OMM), and Loving Kindness Meditation (LKM). Within each graph, colored bands represent the frequency ranges (from left to right): delta, theta, alpha, beta. Significance of differences (*p*) between the two groups is presented in the lower right corner.

As further demonstrated in Figure 4, GC was significantly larger in LTM relative to STM in theta, alpha and beta bands for all connections transferring information from posterior (C3 and C4) to anterior (C1 and C2) clusters. Hence, the increased posterior → anterior information flow in LTM detected also in the time domain (Fig. 2 and 3) was supported by alpha, theta and beta frequencies, with alpha networks being most substantially involved. Of note, the increased information flow across anterior clusters found in LTM in the time domain appears to additionally rely on the strong recruitment of beta networks.

Figure 4 also demonstrates that the increased information transfer across posterior regions observed in the time domain in STM (Fig. 2) was supported by delta networks as indexed by the significantly higher GC in the delta range in STM than LTM for C3 → C4 and C4 → C3 connections. In addition, CG of STM was significantly larger in the delta range for the across-anterior clusters indicating a predominant involvement of delta networks also for directed information flow in frontal regions.

Meditation States: Figure 4 shows that the same between-groups differences in directed information transfer were observed during all meditation states – FAM, OMM, and LKM.

## 4. DISCUSSION

In the present study, Granger causality was analyzed in the time and frequency domains to characterize directed information transfer in highly experienced meditators (LTM) and unexperienced meditators with limited meditation practice (STM). The aim was to reveal effects of long-term meditation on the directed functional connectivity between key cortical regions (frontal and parietal) in the two hemispheres. The second question addressed was if meditation practice affects frequency-specific oscillatory networks supporting directed information flow. To differentiate between neuroplasticity and state-related effects, experienced and unexperienced meditators were compared in a state of rest and three different states of meditation – FAM, OMM and LKM.

Analysis of GC in the time domain revealed that long-term meditation essentially modifies directed information transfer. The major effect was characterized by pronounced increase of information flow (1) from posterior to frontal cortical regions, and (2) across frontal regions of the two hemispheres. Analysis of GC in the frequency domain showed that these dominant transfers were supported by multi-spectral oscillatory networks involving theta, alpha and beta frequency bands, with most prominent expression of GC alpha peak. This pattern of enhanced information transfer in LTM relative to STM was observed in resting state as well as each meditation state. The similarity of between-group differences across states points to a neuroplastic effect of long-term meditation on directed information flow.

Previously, during resting state in non-meditators, a dominant information flow from parieto-occipital to frontal regions has been observed in the higher-frequency bands (alpha and beta), with a simultaneous flow from frontal to temporal and posterior regions in the theta band (Hillebrand et al., 2016). The same frequency-specific pattern has been confirmed in non-meditators for both resting state and memory maintenance conditions (Wang et al., 2019). Current results from unexperienced meditators are in line with these findings by demonstrating a pronounced GC peak in the alpha band for the posterior-to-anterior transfer, and largest GC values of slow-frequency bands (delta/theta) for the anterior-to-posterior transfer. According to Hillebrand et al. (2016), these directional patterns reflect the functioning of major large-scale networks with crucial nodes in the frontal and posterior cortical regions - the Default Mode Network (DMN) and the executive fronto-parietal cognitive networks. The components of these networks were suggested to form a closed loop, through which information “reverberates” or “circulates” with different frequencies of directed information (Hillebrand et al., 2016; Wang et al., 2019). The strong involvement of occipital regions in the loop was further associated with re-entry in neural systems as a mechanism for integration of brain function (Edelman and Gally, 2013). In this perspective, the present analysis includes topological clusters that encompass the nodes of three main large-scale networks identified to provide a functional link between frontal and parietal cortices – DMN, Dorsal Attention Network (DAN), and the Fronto-Parietal Control Network (FPCN) (Uddin et al., 2019). Hence, these networks may be suggested to contribute to the information flow profile observed here in experienced meditators.

During resting state, the DMN typically manifests a prominent posterior → anterior directed connectivity (Uddin et al., 2009; Hlinka et al., 2010). The DMN activation has been linked to spontaneous cognition (internal mentation) and/or general unconscious low-level attention to the external world (Raichle and Snyder, 2007; Buckner et al., 2008). The posterior → anterior information flow within DMN is associated with a role of posterior midline structures in initiating internal mentation, which is then integrated or interpreted by anterior regions (e.g., in self-referential or evaluative contexts). Multi-spectral EEG frequencies from delta (Lauf et al., 2010), theta (Braboszcz and Delorme, 2011; Scheeringa et al., 2008), alpha (Jann et al., 2009; Mantini et al., 2007; Scheeringa et al., 2012), and beta bands (Brookes et al., 2011) have been associated with DMN functioning.

With regard to a possible DMN involvement, the present observations about a significant augmentation of the posterior→anterior information flow in LTM align with previous reports according to which long-term mediation is associated with stronger functional integrity and autonomy of the posterior DMN nodes (Marzetti et al., 2014). In the same line, Rao et al. (2018) have observed that outputs from the frontal DMN nodes (anterior cingulate and orbitofrontal cortices) are significantly reduced in open monitoring meditation implying a dominance of the opposite (posterior→anterior) flow. Hence, within the suggested closed-loop circulation of information in the DMN (Hillebrand et al., 2016), the present results may point to an altered reverberation in LTM characterized by a suppressed frontal output and a dominant influx from posterior to frontal DMN nodes. This functional modulation may reflect a greater initiation of internal processes, such as self-monitoring, meta-awareness, or non-reactive observation of mental content, all of which are trained by meditation practice (Cahn and Polich, 2006; Malinowski, 2013; Lutz et al., 2008; Raffone et al., 2019).

However, such a suggestion is not convincingly supported by other investigations of DMN in long-term meditation. For example, Shaw and Routray (2018) utilized Directed Transfer Function (DTF) and Partial Directed Coherence (PDC) on EEG data to assess directed connectivity in meditation. Opposite to the present results, they found that experienced meditators exhibited increased top-down influences from frontal to posterior brain regions. Barrós-Loscertales et al. (2021) and Garrison et al. (2014) demonstrated decreased connectivity of the precuneous and posterior parietal regions of DMN in meditation. An fMRI study of Brewer et al. (2011) showed that experienced meditators as compared to controls, had reduced activation in core DMN regions (the medial prefrontal cortex and posterior cingulate cortex) across various meditation practices, but increased functional connectivity between the DMN nodes and cognitive control areas in the prefrontal cortex. A meta-analysis of Shen et al. (2020) confirmed that long-term meditation practice was associated with decreased functional connectivity within the DMN and increased connectivity between the anterior DMN and attentional networks. Together, these previous reports do not provide a clear support to the suggestion that the posterior-to-anterior connectivity profile in LTM observed here can only be attributed to enhanced activation of the posterior DMN.

Instead, task-positive fronto-parietal networks may play a role (Hillebrand et al., 2016; Wang et al., 2019). During tasks with goal-directed attention and top-down control, the DAN (Szczepanski et al., 2013; Gross et al., 2004; Popov et al., 2017) and FPCN (Palva and Palva, 2012; FitzGerald et al., 2017; Sadaghiani and Kleinschmidt, 2016) typically demonstrate a fronto→parietal direction of influence supported by increased theta-, alpha-, or beta-directed connectivity. This directional pattern aligns with the role of the frontal cortex in modulating downstream processing based on task goals. Specifically, theta band connectivity from the medial frontal cortex to various regions plays a key role in inducing control over perceptual lower-level processes, monitoring, and coupling with other cognitive networks (Clayton et al., 2015; Cohen, 2011; Cavanagh and Frank, 2014; He et al., 2007; Daitch et al., 2013). Anterior-to-posterior alpha connectivity is proposed to provide a gating mechanism for attention by inhibiting irrelevant activity through top-down modulation (Palva and Palva, 2007; Sadaghiani et al., 2012; Sadaghiani and Kleinschmidt, 2016). Particularly internal attention has been associated with the alpha frequency band (Bonnefond and Jensen, 2013, 2012; Scheeringa et al., 2009; Frey et al., 2015). The anterior-to-posterior flow within executive networks is consistently observed during both resting state and task conditions (Hillebrand et al., 2016; Wang et al., 2019), although reciprocal transfers are also present.

In the framework of the task-related networks, the present findings of reduced frontal dominance and enhanced posterior→anterior information transfer in LTM implies that posterior sensory and associative areas (e.g., visual, parietal cortex) might inform rather than be directed by frontal executive regions. This points to a reduced top-down control, a greater bottom-up integration, and a more receptive or integrative mental state, suggesting a substantial reorganization of attentional and cognitive control networks. It has been acknowledged that posterior regions play a major role for bottom-up attentional capture (Cabeza et al., 2008, 2012) by supporting “cross-modal and multi-modal integrative hubs”, which combine bottom-up inputs with top-down controlling signals (Humphreys et al., 2017) to enhance awareness, manipulate mental events, and reorient attention to relevant information (Seghier, 2013). Similarly, the parietal areas also are associated with the formation of attentional “priority maps” which represent environmental bottom-up features dynamically selected by the top-down focus (Gottlieb et al., 2009). In view of these models, the present results suggest that in LTM, a facilitated formation of strong integrative hubs and stable priority maps in posterior regions may play a leading role in executive control, thus minimizing the contribution of a continuous frontal guidance. Such a reorganization of executive networks may provide neural grounds for the faculty of experienced meditators to sustain a stable and effortless focus of internal attention, monitoring and awareness (Lutz et al., 2008; Raffone et al., 2019).

Current observations of increased bi-directional information flow across the anterior clusters in LTM aligns with previously detected increased connectivity within the anterior DMN, and between anterior DMN nodes and frontal regions of executive networks in relation to meditation (Brewer et al., 2011; Xue et al., 2014; Jang et al., 2011; Taren et al., 2017; Shen et al., 2020). They further imply that in LTM, frontal network coupling is supported by theta, alpha and beta networks. It is to be noted that for all conditions, the outflux from the left frontal cluster was more strongly pronounced than that from the right one, which corresponds to previously observed network modulation and reorganization in the left hemisphere as a function of meditation practice (Marzetti et al., 2014; Fox et al., 2014; Raffone et al., 2019; Yordanova et al., 2020, 2021; De Filippi et al., 2022). It is also interesting to note that in contrast to LTM, the profile of inter-hemispheric directed connectivity in STM was only evident for the posterior clusters and was supported by slow-frequency delta networks (Dissanayaka et al., 2015). In the STM, GC peaked in the delta frequency band (1–4 Hz) also in both anterior-to-posterior and posterior-to-anterior directions, consistent with prior literature linking delta-band oscillations to intrinsic, low-arousal cortical dynamics and DMN activity (Laufs et al., 2003; Cahn and Polich, 2006; Neuner et al., 2014). The dominance of delta-band directed connectivity in STM may therefore reflect a greater prevalence of mind-wandering and self-referential processing, or reduced executive regulation during rest.

In addition to the possible involvement of large-scale cognitive networks and their interactions, the enhanced directed posterior-to-anterior information flow in LTM may be linked more directly to enhanced bottom-up signaling from sensory and interoceptive regions. The spanning of the dominant alpha frequency to theta and beta bands in the enhanced directed information flow is consistent with reports from the primate visual system demonstrating that feedforward influences from occipital regions are carried by theta-band (∼ 4 Hz) and fast-frequency synchronizations (Bastos et al., 2015). In the perspective of intensified transmission of external sensory and interoceptive input to frontal regions, the strongest involvement of alpha frequency may be associated with the recognized faculty of experienced meditators to precisely filter out or inhibit irrelevant sensory signals (Malinowski et al., 2013; Lutz et al., 2008; Yordanova et al., 2025). This suggestion is consistent with models of “gating by inhibition” (Palva and Palva, 2007), according to which alpha rhythms dynamically regulate sensory flow to support attentional and interoceptive processes in meditation (Yordanova et al., 2025). The suggested augmentation of bottom-up signaling also is fully consistent with faculties of long-term meditators characterized by raised awareness of present-moment experience, distributed and receptive sensory processing which is free of elaborative or habitual influences on sensory input.

An alternative explanation for the observed patterns of directed connectivity in LTM may come from the topological properties of the underlying structural networks. Modelling work has suggested that the most strongly connected aspects in a network (hubs) may trigger a stable phase-lag leading to increased functional connectivity (Tomasi and Volkow, 2011). Brain networks have their strongest hubs in posterior regions (Moon et al., 2015), even more so in LTM (Marzetti et al., 2014; Yordanova et al., 2020). It has been emphasized, however, that patterns of phase differences should not be conflated with patterns of directed connectivity (Pullon et al., 2020), and the global strength of functional connections may be accompanied by different directionality profiles (Hillebrand et al., 2016). Instead, the net outflow of information from posterior regions as observed here should be considered as reflecting an increase in encoded information transfer.

Collectively, the present results suggest that long-term meditation practice is associated with a shift in brain dynamics toward reduced reliance on slow, undirected intrinsic oscillations, and enhanced directional connectivity in frequencies linked to attention and cognitive control networks. In addition, the dominant posterior-to-anterior directionality points to a reorganization of cognitive control networks that may support the phenomenological qualities established in long-term meditation — namely, sustained internal attention, enhanced present-moment awareness, and reduced cognitive elaboration. The presence of this dominant directionality pattern during both resting state and different meditative states in LTM highlights adaptive neuroplasticity in large-scale brain networks as a critical outcome of extensive meditation practice.

## Acknowledgments

We would like to express our gratitude and appreciation to the monks, nuns and novices of Amaravati and Santacittarama Buddhist Monasteries for their outstanding dedication and participation in our study. This work has been supported by a grant from BIAL Foundation (Portugal) for the project “Advancements on the aware mind-brain: New insights about the neural correlates of meditation states and traits” (Grant 272/70), and by the National Research Fund by the Ministry of Education and Science, Sofia, Bulgaria (Project KP-06-N33/11/2019).

## Data availability

The datasets used and analyzed during the current study are available from the corresponding author on reasonable request.

